# SARS-CoV-2 protein subunit vaccination elicits potent neutralizing antibody responses

**DOI:** 10.1101/2020.07.31.228486

**Authors:** Marco Mandolesi, Daniel J. Sheward, Leo Hanke, Junjie Ma, Pradeepa Pushparaj, Laura Perez Vidakovics, Changil Kim, Karin Loré, Xaquin Castro Dopico, Jonathan M. Coquet, Gerald McInerney, Gunilla B. Karlsson Hedestam, Ben Murrell

**Author notes:** These authors contributed equally (Marco Mandolesi, Daniel J. Sheward). These authors contributed equally (Gunilla B. Karlsson Hedestam, Ben Murrell).

## Abstract

The outbreak and spread of SARS-CoV-2 (Severe Acute Respiratory Syndrome coronavirus 2), the cause of coronavirus disease 2019 (COVID-19), is a current global health emergency and a prophylactic vaccine is needed urgently. The spike glycoprotein of SARS-CoV-2 mediates entry into host cells, and thus is a target for neutralizing antibodies and vaccine design. Here we show that adjuvanted protein immunization with SARS-CoV-2 spike trimers, stabilized in prefusion conformation ^1^, results in potent antibody responses in mice and rhesus macaques with neutralizing antibody titers orders of magnitude greater than those typically measured in serum from SARS-CoV-2 seropositive humans. Neutralizing antibody responses were observed after a single dose, with exceptionally high titers achieved after boosting. Furthermore, neutralizing antibody titers elicited by a dose-sparing regimen in mice were similar to those obtained from a high dose regimen. Taken together, these data strongly support the development of adjuvanted SARS-CoV-2 prefusion-stabilized spike protein subunit vaccines.

## Introduction

As of 22nd July 2020, at least 15 million cases of SARS-CoV-2 infection have been confirmed, with over 615,000 COVID-19 related deaths recorded ^2^. Cases continue to spread globally despite unprecedented public health measures and lockdowns. An effective prophylactic vaccine is urgently required. Adjuvanted recombinant protein subunit vaccines have excellent safety profiles and represent a proven vaccine platform for eliciting protective immune responses to viral infections, including human papillomavirus (HPV), hepatitis B virus (HBV), and influenza A virus (IAV).

The spike glycoprotein of SARS-CoV-2 mediates receptor binding and entry into target cells, and is the primary target for vaccine design. The receptor binding domain (RBD) is a stable subunit within the spike glycoprotein (Fig. 1A) responsible for ACE2 binding ^3–7^, that can be expressed as an independent domain ^7–9^. While the RBD is a major target for neutralizing antibodies ^10–12^, antibodies specific for spike epitopes outside of the RBD are also capable of neutralization ^13,14^.

**Fig. 1.**
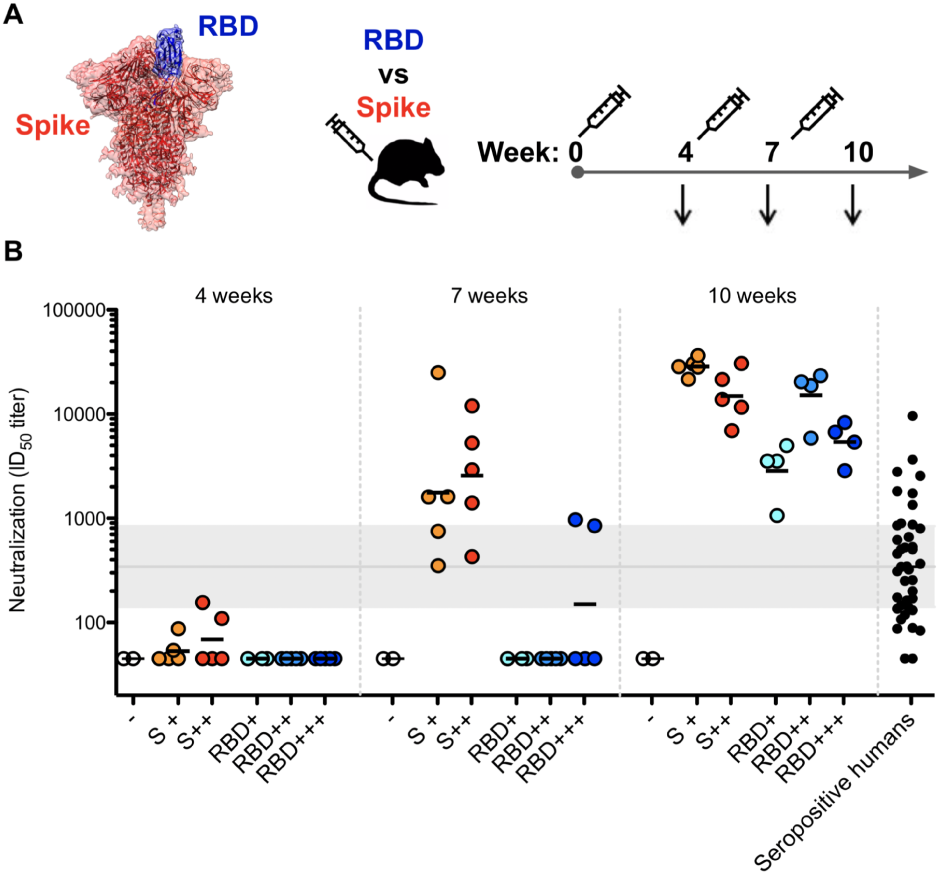
Protein subunit vaccines elicit potent neutralizing antibodies in mice. **A**. Two SARS-CoV-2 protein immunogens were evaluated: stabilized spike trimer, and receptor binding domain (RBD). Mice (N=24) were immunized and humoral immune responses were followed longitudinally to compare stabilized spike vs RBD immunogens at several doses. **B**. Serial dilutions of serum from immunized mice were assessed for neutralization of SARS-CoV-2 pseudotyped lentiviruses harbouring a luciferase reporter gene, and the ID50 titers were calculated as the reciprocal dilution where infection (RLU) was reduced by 50% relative to infection in the absence of serum. The geometric mean ID50 for each group is displayed. - (unimmunized mice - open circles); S+ (5µg stabilized spike - orange); S++ (25µg stabilized spike - red); RBD+ (5µg RBD - cyan); RBD++ (25µg RBD - blue); RBD+++ (50µg RBD - navy). ID50 titers below the limit of detection (45 or 90 depending on sample availability) are displayed as 45. ID50 titers in seropositive donors (black) in Castro Dopico et al. ^18^ determined using the same assay, and the median and interquartile range is highlighted in grey across the background.

## Results

To evaluate the use and immunogenicity of recombinant protein subunit vaccines for SARS-CoV-2 we immunized C57BL/6J mice (N=24) with either the spike ectodomain or RBD, expressed in 293-F cells. The RBD domain was expressed as an Fc-fusion protein, which was cleaved and the RBD subsequently purified by size-exclusion chromatography. The spike ectodomain was expressed as a prefusion-stabilized variant with a C-terminal T4 trimerization domain, and the introduction of two stabilizing Proline mutations in the C-terminal S2 fusion machinery ^1,15,16^. Highly pure and homogenous spike glycoprotein trimers were obtained by affinity purification followed by size-exclusion chromatography (Extended Data Fig. 1). Cryo-EM analysis of the spike preparation demonstrated that it was well-folded and maintained in the trimeric, prefusion conformation^17^, consistent with the original report^1^.

Mice were immunized with varying doses of antigen (range: 5 µg - 50 µg) in Addavax (InvivoGen), a squalene-based oil-in-water emulsion analogous to MF59. MF59 is licensed, safe and effective in humans ^19^, and increases the immunogenicity of an influenza vaccine in the elderly ^20^. Mice were boosted twice, at 3-week intervals, beginning 4 weeks after prime (Fig. 1A). In both the low-dose (5 µg) and high-dose (25 µg) groups, a single immunization with prefusion-stabilized spike elicited a strong spike-specific IgG antibody response, detected by ELISA (Extended Data Fig. 2), as early as 4 weeks following the first immunization. RBD was less immunogenic, with weak to no detectable responses after the initial prime. However, seroconversion was evident in all mice after the first boost with RBD.

To determine whether the antibody responses elicited were neutralizing, we used a SARS-CoV-2 pseudotyped virus neutralization assay. In spike-immunized mice, neutralizing responses (ID50 100) were already detectable in four of ten mice after a single dose (Fig. 1B and Extended Data Fig. 3). All spike-immunized mice developed potent neutralizing antibody responses (median ID50 1,600) after the first boost, which further increased in potency (median ID50 25,000) after the second boost. In contrast, RBD-immunized mice developed consistent neutralizing antibody responses only after the second boost, with a median ID50 neutralizing antibody titer across all groups of 5,300. The substantial enhancement of the virus neutralizing activity observed after the second RBD boost warrants further investigation. Across all groups, spike-specific IgG titers correlated strongly with pseudovirus neutralization, though neutralization was detectable only above a threshold EC50 (Extended Data Fig 5A). Interestingly, RBD medium (25 µg) and high (50 µg) doses elicited neutralizing antibody responses with similar ID50 values to those from the spike groups, but with a significantly flatter slope (p=0.0002), such that they were less neutralizing at higher concentrations (see Extended Data Fig. 4). This could potentially be a result of epitopes on soluble RBD being subject to competition in the context of intact spike trimer. This also suggests an additional advantage of spike immunization over RBD alone, as incomplete neutralization may compromise sterilizing immunity.

Next, we immunized three rhesus macaques (Macaca mulatta) with adjuvanted trimeric prefusion-stabilized spike glycoprotein over approximately 4-week intervals (Fig. 2A), and characterized the titers and kinetics of binding and neutralizing antibodies. Macaques were immunized intramuscularly with 100 µg of spike protein in 75 µg Matrix-M, a saponin-based adjuvant developed for clinical use ^21^ (Novavax AB). Neutralizing antibody responses were already detectable 2 weeks after a single dose (ID50 titers ranging from 150-420) with titers retained at 4 weeks (Fig. 2C). Two weeks after a first boost, the neutralizing antibody responses were extremely potent, exceeding ID50 titers of 20,000 in all three macaques. An additional boost 3 weeks later did not raise the neutralization potency above that obtained with just two immunizations, suggesting a third dose is not needed to achieve highly potent neutralizing antibody responses following spike protein-based vaccination.

**Fig. 2.**
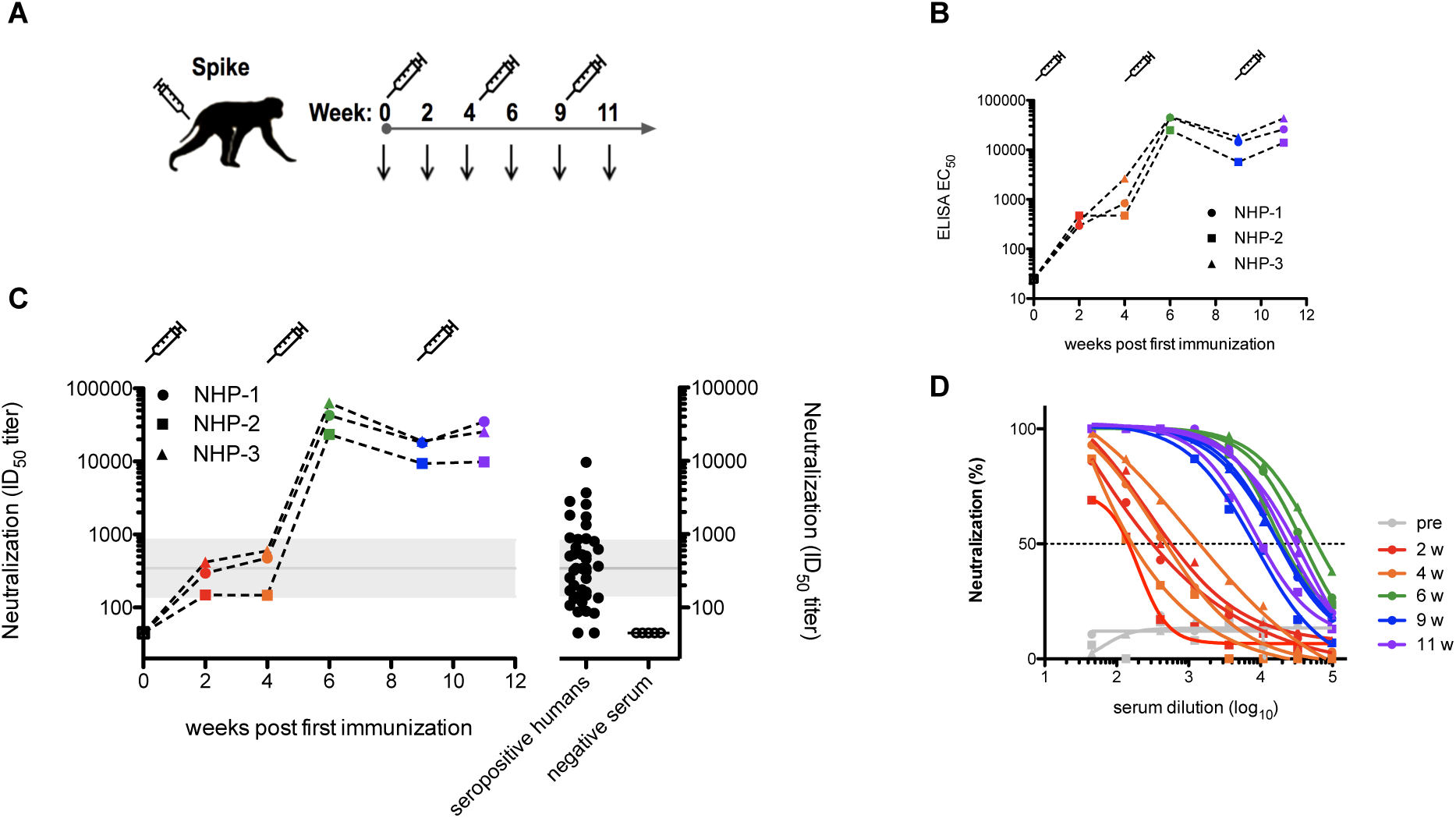
Prefusion-stabilized SARS-CoV-2 spike protein subunit vaccine reproducibly elicits potent neutralizing antibody responses in non-human primates. **A**. Macaque (N=3) immunization and sampling schedule. Syringes indicate the timing of immunizations, and arrows denote times at which blood was drawn. **B**. Longitudinal spike directed IgG responses. **C**. Longitudinal neutralization potency over the course of the study. Neutralization below the assay limit of detection (45) is plotted as 45. Shaded band corresponds to the median and interquartile range of the neutralization potency observed in seropositive human donors using the same assay ^18^, shown on the right. **D**. Neutralization curves depicting percent neutralization against serum dilution.

After two prefusion-stabilized spike protein immunizations, geometric mean neutralizing antibody titers in macaques were approximately two orders of magnitude more potent than that of sera from SARS-CoV-2 seropositive humans (Fig. 2C). These were also substantially higher than those elicited by other immunization platforms that afforded macaques partial or complete protection from challenge^22,23^. In three recent phase I human trials, neutralizing antibody titers elicited by mRNA or adenovirus-vectored vaccines were similar to ^24,25^ or well below ^26^ those from convalescent sera.

The spike-directed IgG binding titers elicited in the macaques (Fig. 2B) correlated strongly with the virus neutralizing activity (R2 = 0.9478, Extended Data Fig. 5) and the rapid development of potent virus neutralizing antibody responses in macaques after a single dose suggested a minimal requirement for somatic hypermutation (SHM) of the response. This is consistent with recent data showing that neutralizing monoclonal antibodies isolated from SARS CoV-2 convalescent individuals display low levels of SHM^12,27,28^. The polyclonality, antibody germline VDJ usage, and level of SHM that characterizes vaccine-induced SARS CoV-2 spike-induced antibody responses will be a matter of interest, both in the macaque model and in human vaccine trials.

## Discussion

Neutralizing antibody titers have varied substantially across different vaccine platforms. In animal models, inactivated virus, DNA, and adenovirus-vectored vaccines elicited neutralizing antibody titers similar to or lower than those seen in convalescent sera ^22,23,29^. While high dose mRNA-based immunizations elicited potent neutralizing antibody responses in mice ^30^, neutralizing antibody titers elicited in phase I human trials were markedly lower^25^. Immunization of mice with mRNA encoding stabilized spike led to reduced or undetectable viral loads in the lungs and nasal turbinates, in a dose-dependent manner, and susceptible transgenic mice were protected from lethal challenge ^30^. Other preclinical vaccine studies using genetic platforms elicited immune responses that protect against disease but not against infection. For example, RBD- and spike-encoding DNA vaccines led to reduced viral loads in the nose and lungs of challenged macaques ^22,23^.

Titers of neutralizing antibodies correlate strongly with protection in a number of vaccine settings ^31–33^. For SARS-CoV-2, passive transfer of neutralizing monoclonal antibodies provided partial or complete protection of animal models, in a dose-dependent manner ^11,34,35^. Here, we show in mice and macaques that neutralizing antibody titers elicited by both spike and RBD protein subunit immunization were orders of magnitude higher than those we detected in SARS-CoV-2 seropositive humans. This reinforces that vaccination can induce more reliable immunity in the population than natural infection. Since it is expected that binding and neutralizing antibody titers decrease from their peak with time, high initial titers will most likely translate into longer lasting protection. At least 19 candidate SARS-CoV-2 vaccines have rapidly entered clinical trials ^36^. These include DNA and RNA-based platforms (Moderna ^25^, Inovio, BioNTech/Pfizer ^37^), adenovirus vectored vaccines (CanSino^26^, University of Oxford/AstraZeneca ^24^, Johnson & Johnson), inactivated virus vaccines (Sinovac, Wuhan Institute of Biological Products, Beijing Institute of Biological Products), and recombinant protein vaccines (Novavax ^38^, Anhui Zhifei Longcom Biopharmaceutical, Clover Biopharmaceuticals/GlaxoSmithKline). Given the urgency in the current pandemic, vaccines will require rapid mobilization on a global scale. As a result, safety is an important consideration, and the use of proven vaccine platforms (particularly in the elderly) would be advantageous. DNA and RNA vaccines are amenable to large-scale and rapid vaccine manufacture; however, no nucleic acid or recombinant adenovirus vaccines are approved for human use, and therefore remain largely untested at scale.

Recombinant protein subunit vaccines have been developed successfully for Influenza, HPV, and HBV, and are also being explored for SARS-CoV-2 vaccination, including three candidate vaccines that have already entered phase I clinical trials. Here we show in animal models (mice and macaques) that immunization with RBD or prefusion-stabilized trimeric SARS-CoV-2 spike proteins elicits potent neutralizing antibody responses. Further, we find that neutralizing responses are more readily elicited and more potent with the stabilized trimeric spike glycoprotein than with RBD. Importantly, we show that neutralizing antibodies were elicited across animal models, using different immunization routes and with different adjuvants, highlighting that the SARS-CoV-2 spike protein represents a robust immunogen. Importantly, significant neutralizing antibody titers arose in non-human primates after a single dose with an adjuvanted spike subunit protein vaccine.

Production of prefusion-stabilized spike at an industrial scale to meet global demand in a rapid time frame may be challenging. However, a number of groups have recently identified additional stabilizing mutations and protocols that dramatically improve protein stability and expression yields ^39–41^. Further, we show in mice that a dose-sparing regimen with 5-fold less spike antigen elicited neutralizing antibodies at a similar titer. The potential to leverage existing infrastructure and manufacturing capacity for licensed vaccines could also accelerate production and rollout.

Here we report that mice and macaques immunized with adjuvanted prefusion-stabilized spike developed potent neutralizing antibody responses. These are stronger than those previously reported to confer partial or complete protection in animal models, and substantially more potent than those from three recent phase I human trials of mRNA or adenovirus-vectored vaccines ^24–26^. Taken together, these results support the development of protein subunit vaccines to immunize against SARS-CoV-2.

## ACKNOWLEDGEMENTS

We thank Dr. Bengt Eriksson and all personnel at Astrid Fagraeus laboratory for expert assistance with rhesus macaques. We also thank Novavax, AB, Uppsala, Sweden, for generously making the Matrix-M adjuvant available. We additionally thank James Voss and Deli Huang for reagents. This project has received funding from the European Union’s Horizon 2020 research and innovation programme under grant agreement No. 101003653 (CoroNAb), to GBKH, GM, and BM, and from the Swedish Research Council to GBKH, GM, and BM.

## AUTHOR CONTRIBUTIONS

Conceptualization, DJS, MM, GBKH, BM; Formal Analysis, DJS, MM, GBKH, BM; Investigation, DJS performed the neutralization assays and the mouse ELISAs. JM performed the mouse immunizations and bleeds. MM, PP coordinated the NHP immunizations, processed samples and performed ELISAs; Resources, LH and LPV produced RBD and spike immunogens, CK and DJS produced pseudovirus, XCD contributed data from seropositive humans; Visualization, DJS, LH, BM; Writing – Original Draft, DJS, BM; Writing – Review & Editing, DJS, KL, JC, GM, BM, GBKH; Funding Acquisition, BM, GBKH, GM; Supervision, BM, GBKH, JC, GM.

## METHODS

### Ethics statement

The animal work was conducted with the approval of the regional Ethical Committee on Animal Experiments (Stockholms Norra Djurförsök-setiska Nämnd). All animal procedures were performed according to approved guidelines.

### Protein production

The plasmid for expression of the SARS-CoV-2 prefusion-stabilized spike ectodomain ^1^ was kindly provided by Jason McLellan. This plasmid was used to transiently transfect freestyle 293-F cells using the FreeStyle MAX reagent (Thermo Fisher Scientific). The spike ectodomain was purified from filtered supernatant on Streptactin XT resin (IBA Lifesciences), followed by size-exclusion chromatography on a Superdex 200 in 5 mM Tris pH 8, 200 mM NaCl.

The RBD domain (RVQ-VNF) of SARS-CoV-2 was cloned upstream of an enterokinase cleavage site and a human Fc. This plasmid was used to transiently transfect FreeStyle 293F cells using the FreeStyle MAX reagent. The RBD-Fc fusion was purified from filtered supernatant on Protein G Sepharose (GE Healthcare) and cleaved using bovine enterokinase (GenScript) leaving a FLAG-tag on the C-terminus of the RBD. Enzyme and Fc-portion was removed on His-Pur Ni-NTA resin and Protein G Sepharose respectively, and the RBD was purified by size-exclusion chromatography on a Superdex 200 in 5 mM Tris pH 8, 200 mM NaCl. Proteins were re-buffered into PBS prior to immunization.

### Mice

C57BL/6J (Jackson Laboratory) wild-type mice were housed and bred at Karolinska Institutet animal facility. Immunogens were diluted in sterile PBS, emulsified in AddaVax (InvivoGen) and injected subcutaneously (s.c.) in the flanks of mice. Each arm contained 5 mice, except the low-dose RBD group which had 4. Two control mice were not immunized. Tail bleeds were taken prior to each immunization. Whole blood was allowed to clot at room temperature, and serum was separated by centrifugation, heat inactivated at 56°C for 1 hour, and stored at -20°C until use.

### Macaques

Two female and one male rhesus macaques (Macaca mulatta) of Chinese origin, 4-5 years old, were housed at the Astrid Fagraeus Laboratory at Karolinska Institutet. Housing and care procedures complied with the provisions and general guidelines of the Swedish Board of Agriculture. The facility has been assigned an Animal Welfare Assurance number by the Office of Laboratory Animal Welfare (OLAW) at the National Institutes of Health (NIH). The macaques were housed in groups in 14 m3 enriched cages. They were habituated to the housing conditions for more than six weeks before the start of the experiment and subjected to positive reinforcement training in order to reduce the stress associated with experimental procedures. All immunizations and blood samplings were performed under sedation with 10-15 mg/kg ketamine (Ketaminol 100 mg/ml, Intervet, Sweden) administered intramuscularly (i.m.). The macaques were weighed at each sampling. All animals were confirmed negative for simian immunodeficiency virus (SIV), simian T cell lymphotropic virus, simian retrovirus type D and simian herpes B virus.

For macaque immunizations, stabilized spike trimer (100 µg) was mixed in 75 µg of Matrix-M (Novavax AB). Macaques were immunized intramuscularly (i.m.) with half of the dose administered in each quadricep.

### Mouse ELISAs

ELISA plates (Nunc MaxiSorp, Thermo Fisher Scientific) were coated overnight at 4°C with 100 µl of prefusion-stabilized spike protein at a concentration of 1 µg/ml in 1x PBS. Plates were blocked for 90 minutes at room temperature with 200µl of a blocking solution containing 2%(w/v) non-fat milk powder in 1X PBS, and washed 6 times with 1X PBS supplemented with 0.05% Tween 20 (PBS-T). Serum samples serially diluted in blocking solution were added and plates were incubated overnight at 4°C. Plates were washed 6 times with PBS-T, and 100µl of a goat anti-Mouse IgG horseradish peroxidase (HRP) conjugated secondary antibody (Southern Biotech) diluted 1:5,000 in PBS-T was added to each well. Plates were washed 6 times with PBS-T, developed for 15 minutes at room temperature using 200 µl per well of peroxidase substrate (o-phenylenediamine dihydrochloride, SIGMAFAST, SigmaAldrich), and read at 450 nm in an Asys Expert 96 plate reader (Biochrom). EC50 titers were calculated using a Bayesian logistic curve fitting approach, allowing plate-specific minimum and maximum sigmoid parameters to account for differences between plates, and sample specific slope and offset parameters. EC50 titers were calculated from the posterior median value midway between the plate minimum and maximum.

### Macaque ELISAs

ELISA plates were coated with prefusion-stabilized spike protein as above, washed, and blocked with 5%(w/v) non-fat milk powder in 1x PBS. Thereafter, 10-fold serial dilutions (starting at 1:200) of plasma in blocking buffer were added, and plates were incubated for 2 hours at room temperature. Plates were washed and antibody-antigen interaction was detected using HRP-conjugated anti-monkey IgG Fc (#GAMon/IgG(Fc)/PO Nordic MUbio) diluted 1:20,000 in PBS-T. Plates were developed using 50 µl of 3,3’,5,5"-tetramethylbenzidine substrate solution (Invitrogen) per well and stopped using 50 µl of 1M sulphuric acid per well. OD was read at 450nm in an Asys Expert 96 plate reader (Biochrom). Every washing step was performed with 0.05% PBS-T. All experiments were conducted in triplicates. EC50 titers were calculated as for the mice.

### Pseudotyped neutralization assays

Pseudotyped neutralization assays were adapted from protocols previously validated to characterize the neutralization of HIV ^42^ but with the use of HEK293T-ACE2 cells, as previously described ^17^. All cell lines were cultured in a humidified 37°C incubator (5% CO2) in Dulbecco’s Modified Eagle Medium (Gibco) supplemented with 10% Fetal Bovine Serum and 1% Penicillin/Streptomycin, and were passaged when nearing confluency using 1X Trypsin-EDTA. All cell lines tested negative for mycoplasma by PCR. Pseudotyped lentiviruses displaying the SARS-CoV-2 spike protein (harboring an 18 amino acid truncation of the cytoplasmic tail) and packaging a luciferase reporter gene were generated by the co-transfection of HEK293T cells using Lipofectamine 3000 (Invitrogen) per the manufacturer’s protocols. Media was changed 12-16 hours after transfection, and pseudotyped viruses were harvested at 48- and 72-hours post transfection, filtered through a 0.45 µm filter, and stored at -80°C until use. Pseudotyped viruses sufficient to generate 100,000 RLUs were incubated with serial dilutions of serum for 60 min at 37°C in a 96-well plate, and then 15,000 HEK293T-ACE2 cells were added to each well. Plates were incubated at 37°C for 48 hours, and luminescence was then measured using Bright-Glo (Promega) per the manufacturer’s protocol, on a GM-2000 luminometer (Promega). ID50 titers were calculated as the reciprocal serum dilution at which RLUs were reduced by 50% relative to control wells in the absence of serum.

---

**Fig. ED1.**
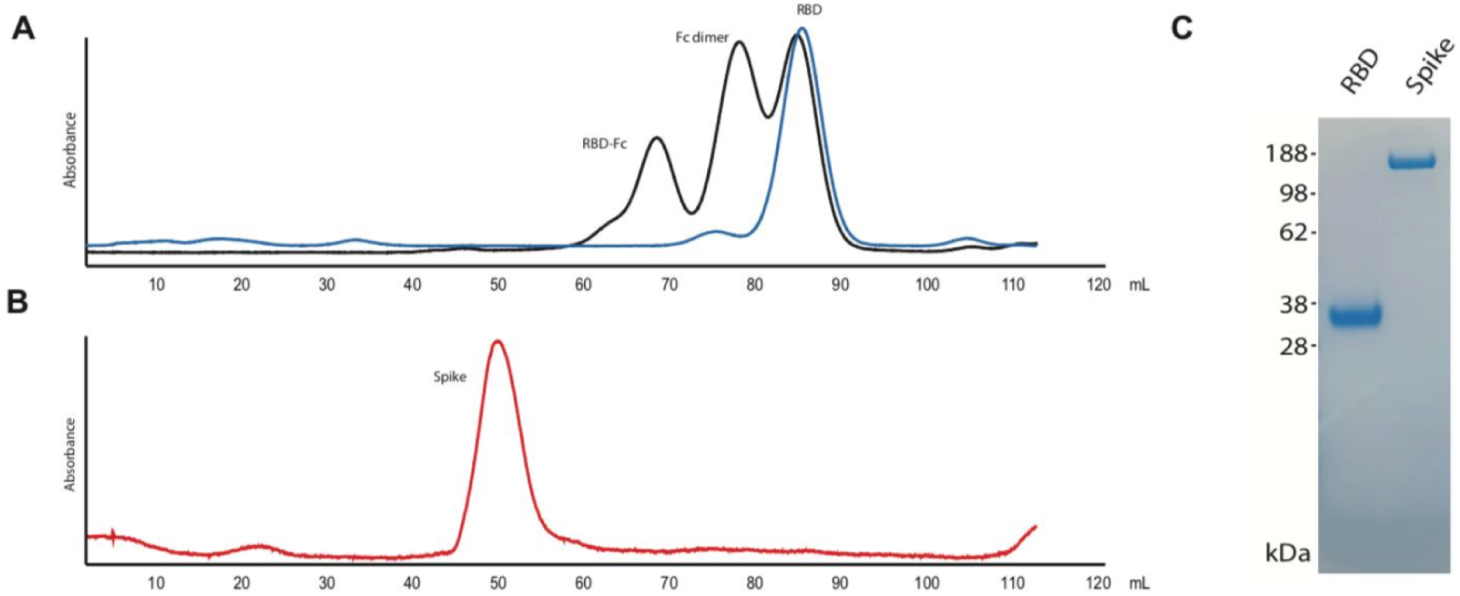
SARS-CoV-2 antigen production. **A**. Size-exclusion chromatograms of affinity purified RBD after enterokinase cleavage digest, before (black) and after (blue) enzyme and Fc removal. **B**. Chromatogram of affinity purified prefusion-stabilized spike ectodomain. Data from a Superdex S200 16/600 **C**. SDS-PAGE analysis of purified SARS-CoV-2 RBD and spike immunogens. Maintenance of spike immunogens in the trimeric, prefusion conformation was confirmed by cryo-EM ^17^.

**Fig. ED2.**
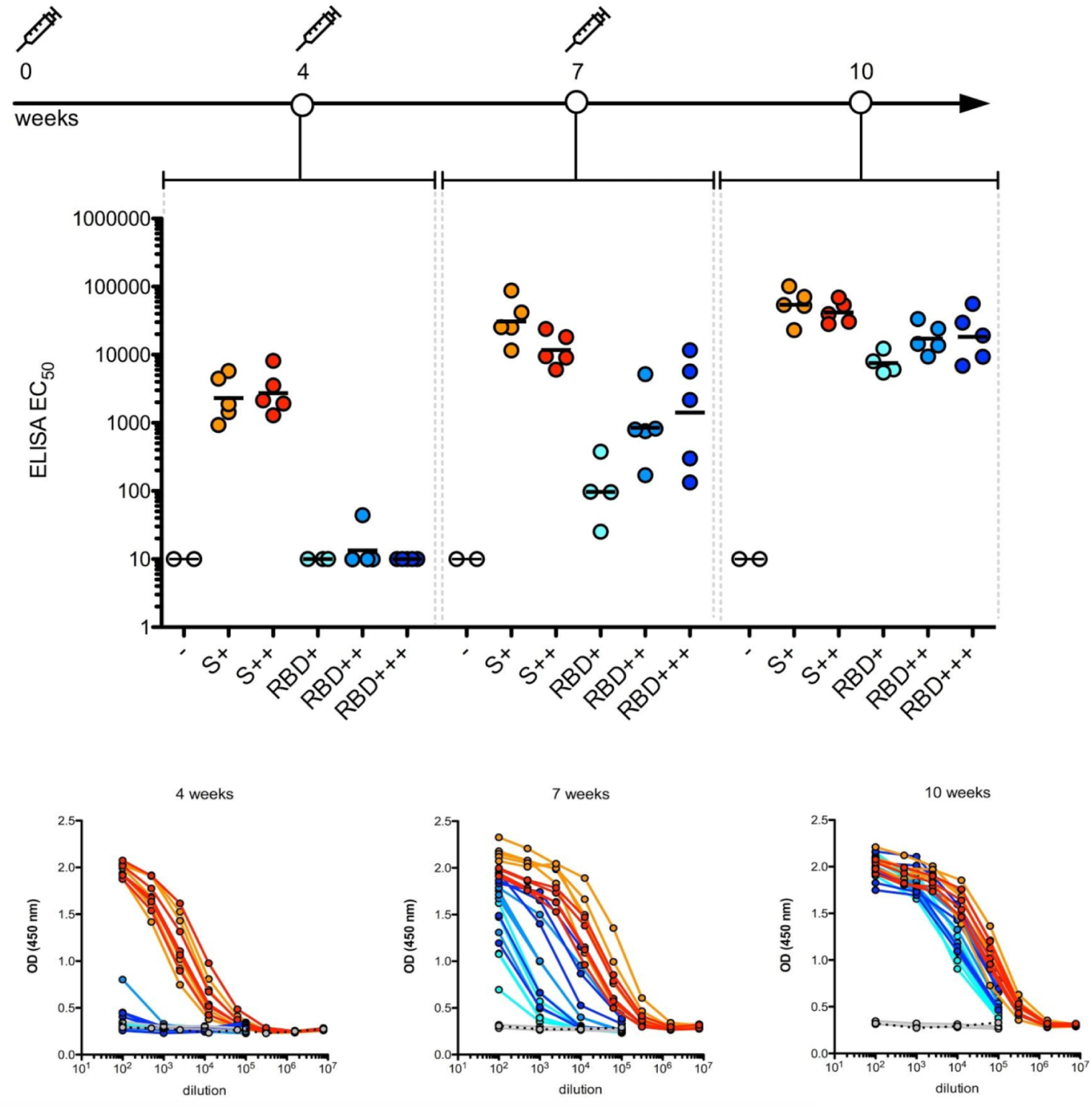
Spike and RBD protein subunits are immunogenic in mice. IgG antibodies detected by ELISA in immunized and control mice reveal that stabilized spike and RBD elicit antibody responses after one, two, and three doses. - (unimmunized - open circles); S+ (5µg stabilized spike - orange); S++ (25µg stabilized spike - red); RBD+ (5µg RBD - cyan); RBD++ (25µg RBD - blue); RBD+++ (50µg RBD - navy).

**Fig. ED3.**
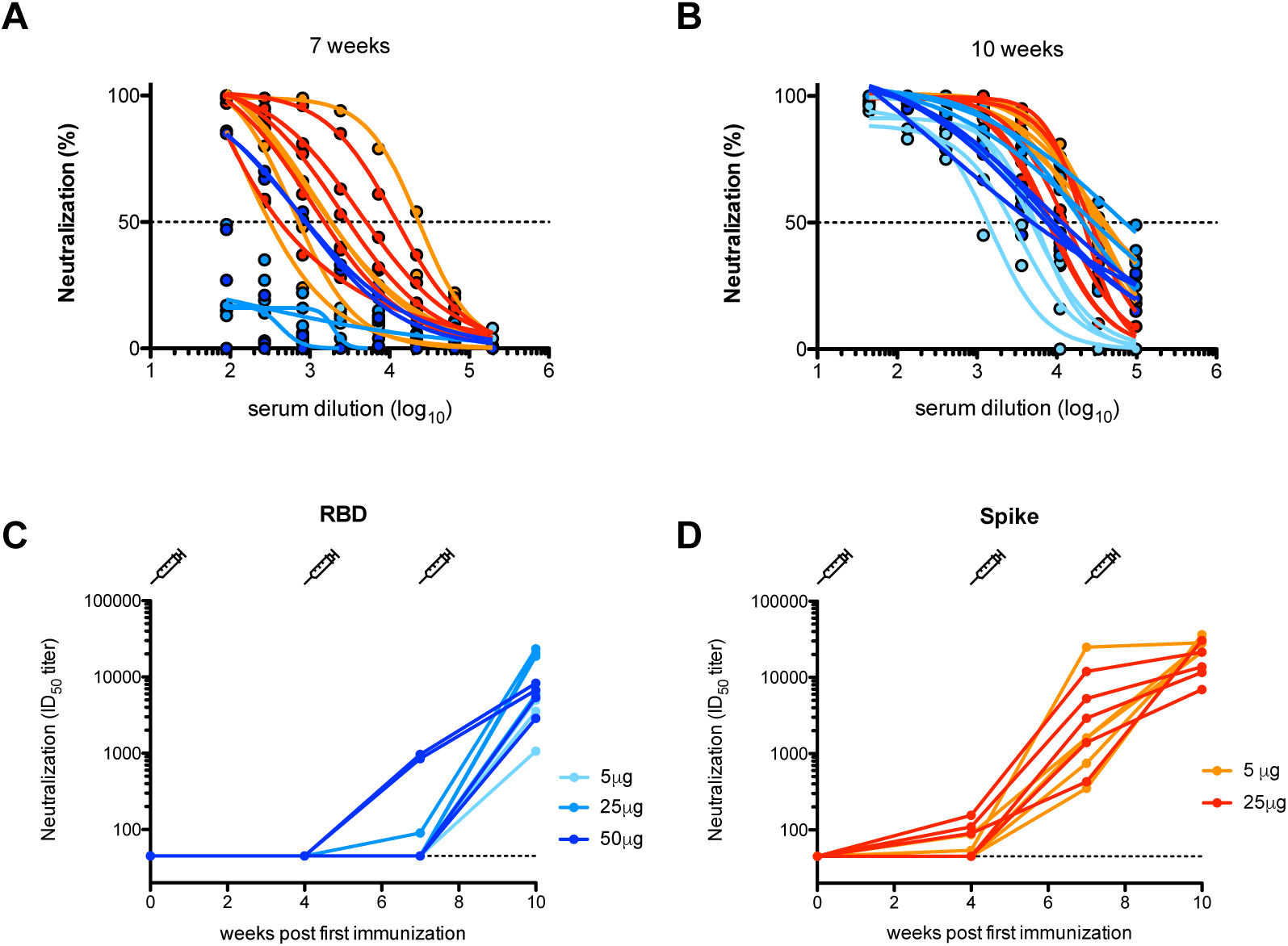
**A-B.** Neutralization curves for mouse sera sampled 7- and 10-weeks post-immunization. Neutralization by serum from mice immunized with RBD are displayed in shades of blue (5µg doses, cyan; 25µg, blue; 50µg, navy). Neutralization by serum from mice immunized with prefusion-conformation stabilized spike ectodomain are displayed in orange (5µg doses) and red (25µg). **C-D**. Longitudinal neutralizing antibody ID50 titers in mice immunized with RBD (**C**) or spike (**D**). Syringes depict the timing of immunizations (0-, 4- and 7-weeks).

**Fig. ED4.**
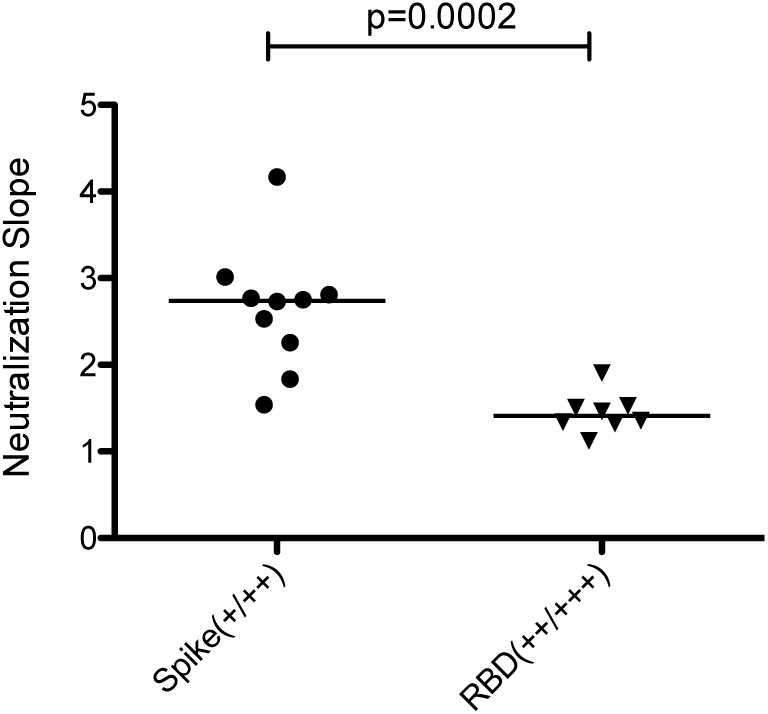
Comparison of the slope of neutralizing antibody curves in 10-week serum (see Extended Data Fig 2B) from Spike or RBD (medium and high dose) immunized mice.

**Fig. ED5.**
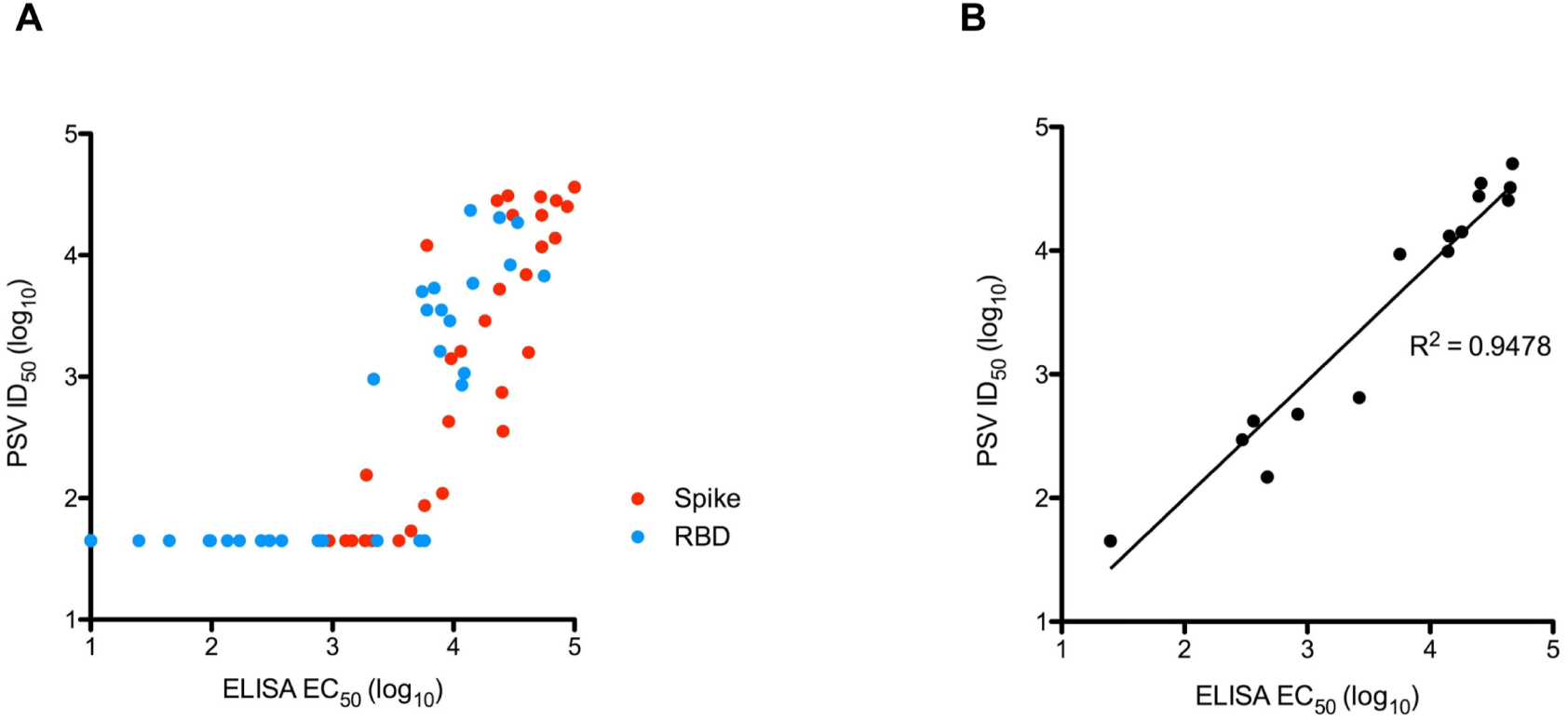
Correlation between spike IgG ELISA EC50 values and pseudovirus (PSV) neutralization ID50 titer in immunized mice (**A**) and macaque (**B**) sera. Neutralization ID50 below the limit of detection is plotted as the assay’s limit of detection (45).

